# Inferring the regulation dynamics of oscillatory networks from scRNA-seq data

**DOI:** 10.1101/2025.11.08.687360

**Authors:** Wenjun Zhao, Alma Plaza-Rodriguez, Pichayathida Luanpaisanon, Elena Xinyi Wang, Linnéa Gyllingberg, Ritambhara Singh, Elana J. Fertig, Genevieve L. Stein-O’Brien

**Affiliations:** Department of Mathematics, Wake Forest University, 2601 Wake Forest Rd, Winston-Salem, NC 27106; Institute for Genome Sciences, University of Maryland School of Medicine, 670 W Baltimore St, Baltimore, MD, 21201, USA; Department of Biomedical Engineering, University of Virginia PO Box 800759, Charlottesville, VA 22908; Department of Informatics, University of Fribourg, Bd de Pèrolles 90, 1700 Fribourg, Switzerland; Department of Mathematics, University of California, Los Angeles, 520 Portola Plaza, Los Angeles, CA, 90095, USA; Department of Computer Science, Center for Computational Molecular Biology, Brown University, 164 Angell St, Providence, RI, 02912, USA; Department of Neuroscience, Johns Hopkins University, 3400 North Charles Street, Baltimore, MD 21218

## Abstract

Oscillatory processes such as the cell cycle play critical roles in cell fate determination and disease development. Yet, most current gene regulatory network (GRN) inference methods are based on gene-gene correlations or temporal progression, not adequately accounting for the recurrence in cyclic processes. We hypothesize that constraining the continuous ordering of relative positions along the cell cycle can enhance GRN inference accuracy of cell cycle regulation. To test performance, we evaluated eight representative methods and applied three of them to a mouse retinal progenitor single-cell gene expression dataset [1]. Incorporating cell cycle positions inferred by Tricycle [2] led to significant improvements compared against using experimental times, particularly for early progenitor cells that been hypothesized to be more intrinsically driven by cell cycle regulation. These findings highlight the promise of integrating oscillatory processes into causal inference frameworks to advance our understanding of gene regulation.

## 1 Introduction

Despite their importance, oscillatory signals are often overlooked in current gene regulatory network (GRN) inference methods because of the inherent difficultly and cost to obtain the measurements required and the inherent lack of coupling between measurements. Current high-throughput gene expression profiling techniques kill the cell they are measuring. Thus, most algorithms focus exclusively on static networks, inferring regulatory relationships from snapshot measurements without incorporating temporal structure (e.g., [3, 4]). While such approaches may recover co-expression patterns, they cannot distinguish direct from indirect interactions or capture causal dynamics, particularly in systems governed by cyclic or transient regulatory programs.

Methods that incorporate temporal information (e.g., [5, 6]) typically rely on experimental time stamps or pseudotemporal orderings inferred from differentiation trajectories, which are fundamentally unidirectional and non-repeating. However, biological processes such as cell cycle progression, Notch cycling [7], and circadian rhythms [8] operate on different time scales and exhibit fundamentally divergent, yet recurrent structures in gene expression space. In these cases, the assumed time axis—whether measured or inferred—may not align with the intrinsic timing of the regulatory process, introducing systematic biases that propagate through the inference pipeline. This misalignment may explain why temporal GRN methods often perform worse than their static counterparts in such contexts [9].

We hypothesize that explicitly modeling the continuous ordering of cyclic temporal processes can improve GRN inference. By embedding cells or samples along a latent trajectory that preserves oscillatory structure, as inferred by Tricycle [2]—a transfer learning method that assigns cycle positions based on prelearned mani-fold—we can better constrain the timing of regulatory events and enhance causal inference. This perspective not only aligns with emerging insights into dynamic gene expression programs but also offers a promising framework for integrating single-cell and time-series data across multiple time scales and contexts.

## 2 Overview on GRN inference algorithms

In this work, we select eight representative methods from the state-of-the-art algorithms, ranging from linear regression-based methods to modern deep learning models, each producing a weighted graph as outcome. We provide a detailed overview of each method as below, and a comparison on the input and output type is provided in Table 1.

**Table 1:** Comparison of gene regulatory network attributes for the methods. We compare the methods based on their capability of (1) distinguishing source and target of causal relations (directed/undirected), (2) predicting whether the type of regulation is activation or repression (signed/unsigned), and (3) integrating temporal information.

| Method | Directed/Undirected | Signed/Unsigned | Temporal |
| --- | --- | --- | --- |
| Scribe | Directed | Unsigned | Yes |
| GeneLink | Undirected | Unsigned | No |
| SINGE | Directed | Unsigned | Yes |
| ARACNE | Undirected | Unsigned | No |
| GENIE3 | Directed | Unsigned | No |
| SINCERITIES | Directed | Signed | Yes |
| SWGCNA | Undirected | Signed | No |
| OTVelo | Directed | Signed | Yes |

### ARACNE

ARACNE [4] is a computational method used to infer gene regulatory networks from gene expression data. It is based on information theory and utilizes mutual information (MI) to identify statistical dependencies between gene expression profiles, which can suggest regulatory relationships. two main steps: 1. Calculate mutual information between all pairs of genes to detect potential regulatory interactions. 2. Apply the Data Processing Inequality (DPI) to eliminate indirect interactions by removing the weakest edge in any triplet of genes that share mutual information, thus reducing false positives. The result is a more accurate and sparse network that reflects direct regulatory relationships between genes, making ARACNE a powerful tool for understanding transcriptional regulation in complex biological systems.

### GENIE3 [3]

GENIE3 is a tree-based method that models the expression level of each gene using all other genes. It was originally designed for bulk data and does not incorporate temporal information. The inferred links are directed based on regressor–outcome pairs, and their importance is ranked according to the variance reduction achieved by including each edge. GENIE3 does not rely on time point information and can work on single time point snapshot data. Notably, it was ranked as one of the top performers in benchmarking challenges [9].

### ODE based method [10]

Ma et al. have developed a method including both time series and steady state data in order to construct GRNs from gene expression data. They model gene interactions using nonlinear ordinary differential equations (ODEs) and then apply the XGBoost algorithm [11] to identify regulatory relationships between genes [10]. The approach aims to combine the dynamic accuracy of ODE-based models with the computational efficiency of machine-learning-based feature selection. While the method shows higher performance than popular approaches such as GENIE3 on the DREAM4 benchmarks, the method relies on highly specific hyper-parameter settings that must be carefully tuned for each DREAM4 dataset, making its performance heavily dependent on dataset-specific optimization and impractical for data without known ground truth.

### SINCERITIES [5]

SINCERITIES is a method that applies Granger causality to the Kolmogorov-Smirnov test statistic on time-stamped data. It assigns the importance of each interaction based on a linear regression of these statistics between consecutive time points. Finally, the signs of the edges are determined using partial correlation.

### SINGE [12]

SINGE is an algorithm that takes ordered single-cell gene expression data. It uses kernel-based regression to smooth out noise in the data and runs over an ensemble of different hyperparameters, such as kernel bandwidth, to prioritize reliable regulatory relationships.

### GENELink and GNNLink

GENELink[13] employs a Graph Attention Network (GAT) to capture the regulatory relationships between transcription factors and target genes. It incorporates both scRNA-seq expression data and prior TF-gene networks to learn low-dimensional gene embeddings. By leveraging multi-head attention mechanisms and MLP channels, it optimizes these embeddings for predicting pairwise regulatory links or causal interactions.

GNNLink [14], in contrast, uses a Graph Convolutional Network (GCN) as its core architecture. It preprocesses scRNA-seq data and constructs an initial GRN from known interactions, then learns gene features through a GCN-based encoder. Final link predictions are generated via matrix completion using the learned node features.

### OTVelo [15]

Given time-stamped data, OTVelo first infers trajectories through optimal transport and approximates the rate of change using finite differences. Granger causality is then applied to infer regulatory interactions between these velocity estimates. It offers two different modalities: an efficient time-lagged correlation approach and a causality approach based on linear regression.

The remainder of the paper will be limited to the 8 methods above. To decide on which methods to use for real data, we first implemented all these methods using the DREAM4 challenge *in silico* gene expression datasets [16] that had a known ground truth. The investigation of usability helped to rule out methods that (1) require careful hyperparameter tuning such as the ODE methods, SINGE, OTVelo, (2) take additional supervising information such as GENELink and GNNLink, and (3) we could not execute during the workshop in which we analyzed these data such as Scribe.

Ultimately, we selected three algorithms that are most user-friendly: (1) GENIE3 [3], which does not use any temporal information and requires downsample to at most 1000 cells due to prohibitive computational cost, (2) ARACNE [4], which is also static and represent edge strengths through mutual information, and (3) SINCERITIES [5], which takes temporal bins and is ideal for testing our hypothesis on the effect of different choices on pseudotime or cell cycle time.

## 3 Results on mouse retinal progenitor cells

While DREAM4 data [16] is useful for containing a known ground truth, the generic stochastic differential equations that define that ground truth do not address the motivation to incorporate oscillatory processes. Thus, we applied the algorithms to an experimental dataset [1] that profiled mouse retinal development using single-cell RNA-seq to investigate whether incorporating oscillatory structure improves performance.

The mouse developing retina is a well characterized system that is often used as a model for more complex developmental processes in the central nervous system, i.e. the cerebral cortex. The entire structure is generated from a single set of progenitors over a stereotyped timeline. As the progenitors move through development, the length of cell cycle increases in a manner that is strongly associated with changes in daughter cell fate specification.

Previous experiments have demonstrated that perturbations to the length of cell cycle will also perturb the type of neurons that the progenitors are able to make at the same developmental age [1]. Additionally, the lengthening of cell cycle is has been linked to restriction in the types of daughter cells which the progenitors are able to generate—a phenomena called progenitor competence. Thus, accounting for cell cycle is critical to inferring the GRN controlling the evolution of progenitor competence and understanding how particular fates are specified during retina development.

To see if we could isolate the shifts in progenitor competence, we applied the GRN inference algorithms on a subset of cells and genes from [1]. Specifically, cells were subset to either early-stage retinal progenitor cells (early RPCs) or late-stage retinal progenitor cells (late RPCs) with both groups genes expression restricted to the same set of 3,164 high variance genes. To further test our hypothesis that incorporating the dynamics of oscillatory processes may improve performance, we provided the algorithms that use temporal information with the following three types of real or pseudo-time orderings:

1. **Real time:** Raw experimental time points, taken at various hours (11, 12, 14, 16, 18, 20, 22, 25, and 28), with most cells concentrated at early time points;
2. **Tricycle time:** Tricycle coordinates obtained using cell cycle genes and the tricycle software [2], which effectively captured the relative position of each cell during the cell cycle. We discretized these into 10 bins based on quantiles;
3. **Integrated** *θ* **time:** A new time coordinate that integrated real time and tricycle time according to a weighted sum, such that each consecutive pair of time points was separated by 2*π*, with their relative magnitudes determined using tricycle time. We discretized this into 10 bins based on quantiles.

The resulting weighted network is benchmarked through comparison with cell type-specific interactions from databases. We inferred gene signatures associated with all RPCs as well as specific to early and late RPCs through CoGAPS analysis [17] with 80 patterns of our scRNA-seq atlas of retinal development as described previously in Clark et al [1] and Stein-O’Brien et al [18]. Sets of genes were associated with all RPCs, early RPCs, and late RPCs by applying the CoGAPS pattern marker statistic [17] to infer unique genes associated with each pattern. Early and late RPC networks were formed by merging the all RPC gene set with the early and late gene sets, respectively, and formulating networks of molecular interactions with STRING database [19]. The pattern marker and network files are available as supplemental files.

### 3.1 Tricycle coordinates improve accuracy metrics

The network structure from STRING is derived from independent experiments of protein-protein interaction for key nodes inferred in our retinal datasets, enabling its use as a ground truth against which to benchmark entirely data-driven networks. Since the ground truth subnetwork was quite sparse (density equals 0.03 for early RPC and 0.08 for late RPC), we report a normalized version of the area under the precision-recall curve (AUPR), defined as the area divided by the network density, following the metric suggestions from BEELINE [9]:

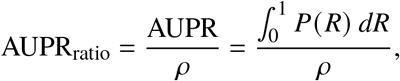

where 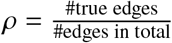 is the density of ground truth network, *P*(*R*) is precision as a function of *R*, and 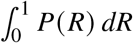 is the area under precision-recall curve, a canonical metric for assessing binary classification accuracy. Under random edge assignment, the expected AUPR equals the density *ρ*, so the AUPR ratio equals 1, providing a convenient normalized baseline.

The results are shown in the barplot in Figure 1. As expected, for the algorithm using temporal information (SINCERITIES), incorporating cell-cycle-based time information significantly improved the accuracy over actual time across the board. In particular, incorporating cell cycle-based time yielded the best performance of all metrics on early RPCs reflecting the experimental evidence that early RPCs are more intrinsically driven by cell cycle regulation.

**Fig. 1:**
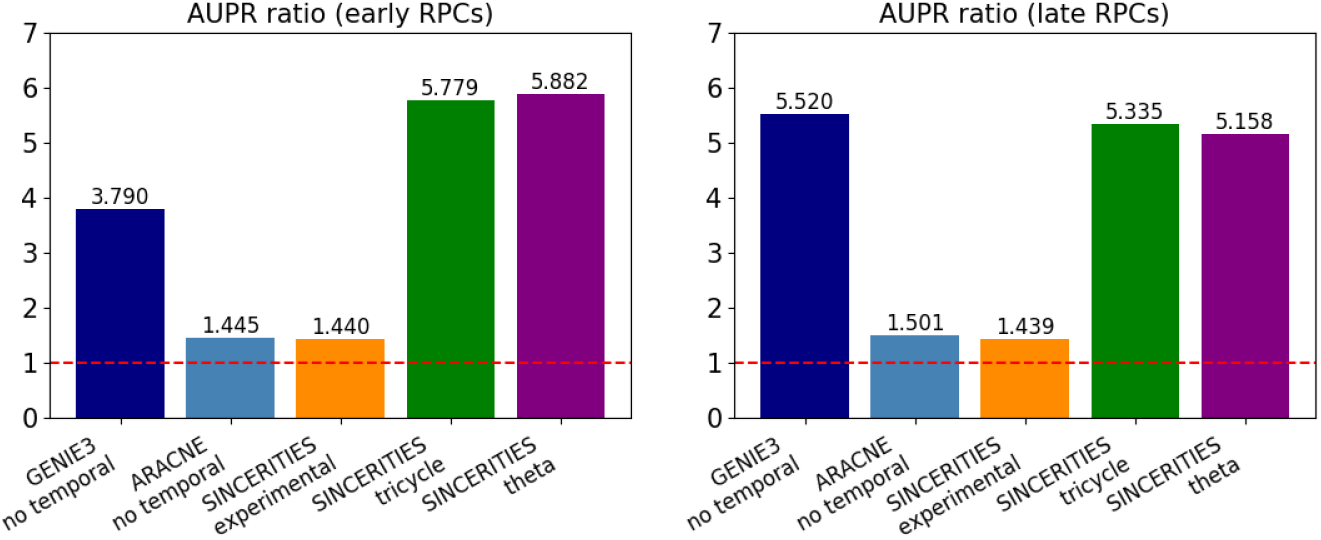
AUPR ratio values for early RPCs (left) and late RPCs (right). The baseline of a random classifier is 1.0, indicated by the horizontal red dashed line.

While incorporating cell cycle-based time information also significantly improved the accuracy over actual time in late RPCs, the increased performance of GENIE3 which relies on regressor–outcome pairs suggests signaling networks in addition to cell cycle playing a role in determine competence of late RPCs. These findings support the hypothesis that progenitor competence of late RPCs is more extrinsically driven and that the lengthening of cell cycle is there to allow the cells additional time to process signals from their environment.

For early RPCs, we hypothesize that the improvement in performance is likely due to better incorporation of cell cycle gene information in the tricycle and theta time orderings. In particular, we examined two cell cycle genes, *Top2a* and *Ccnb1*, to assess how the inferred networks from early RPCs captured their interactions. For simplicity, we visualized the top 10 edges related to *Top2a* and *Ccnb1* in Table 2, and compared them to the STRING network shown in Figure 2. As a result, all edges inferred by SINCERITIES using theta time were supported by the STRING network. SINCERITIES using tricycle coordinates had only 1 out of 20 edges that is not supported. GENIE3 had 80% supported edges, while only one edge was supported when SINCERITIES used the actual experimental time. ARACNE infers 50% correct edges for *Top2a*, and misses all edges for *Ccnb1*.

**Table 2:** Top 10 edges involving *Ccnb1* and *Top2a* from the inferred networks for early RPCs. Dark blue edges indicate validation by STRING (Figure 2), while light blue indicates unsupported edges.

| Algorithm | Ccnb1 | Top2a |
| --- | --- | --- |
| GENIE3 (no temporal) |  |  |
| ARACNE (no temporal) |  |  |
| SINCERITIES (experimental) |  |  |
| SINCERITIES (tricycle) |  |  |
| SINCERITIES (theta) |  |  |

**Fig. 2:**
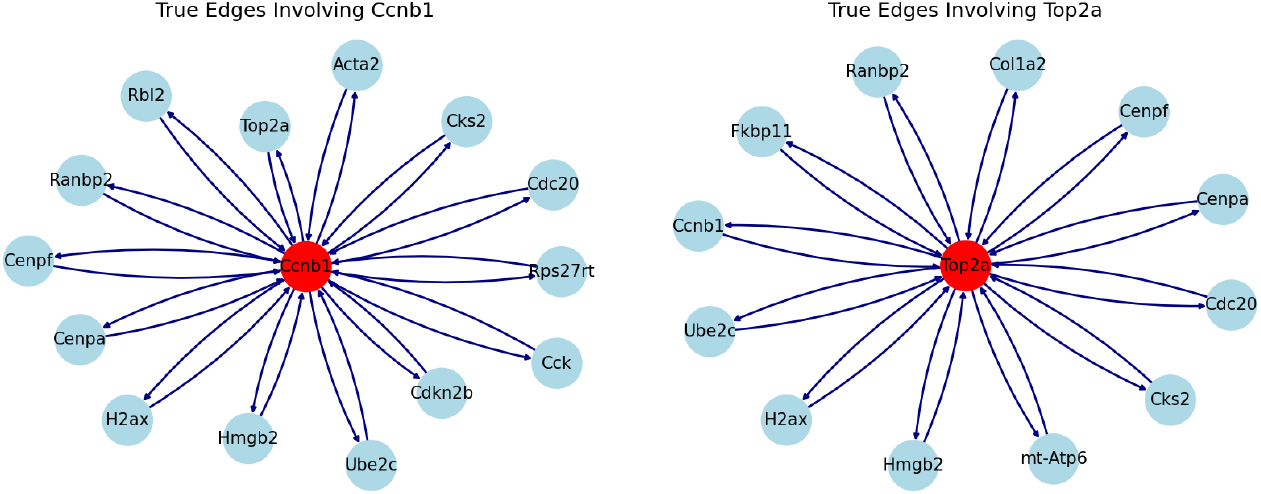
STRING network involving *Top2a* and *Ccnb1*. These two genes were selected because (1) they are involved in the cell cycle, and (2) they have more known interactions according to the database.

### 3.2 Tricycle coordinates identify hub genes relevant for biological processes

GRN inference algorithms using mouse retinal development data yielded complex networks. Our goal was to prioritize the top hub genes for both early- and late-stage progenitor cells. Centrality analyses were performed on the final inferred networks to quantify centrality of degree, closeness, and betweeness, as measures to identify hub genes. We focused on degree of centrality, which prioritizes top genes across different methods. The centrality is defined as the sum of all incoming importance for each target gene. The results for early and late RPCs, across 5 algorithms, are shown in Table 3.

**Table 3:**
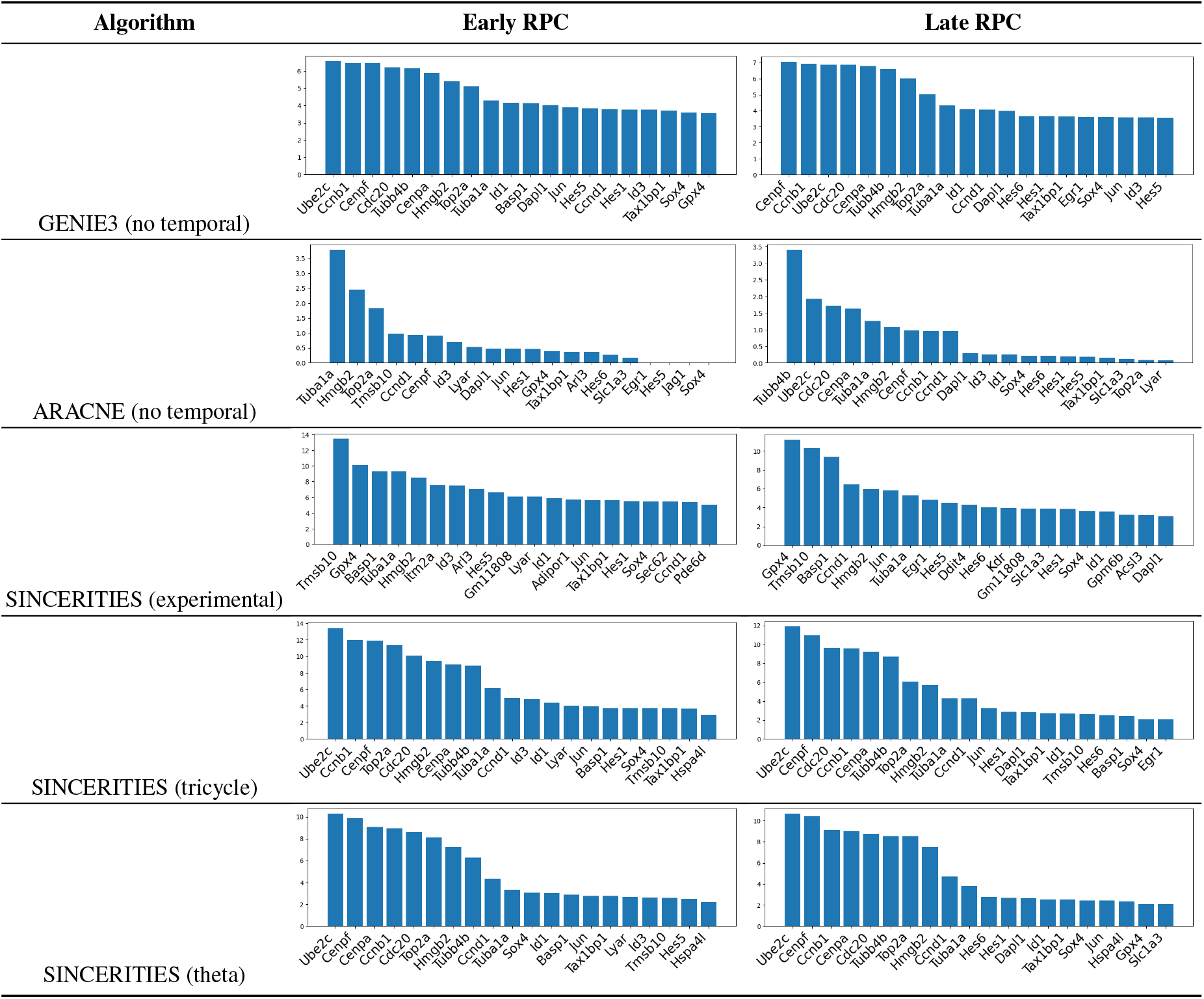
Comparison of hub genes identified by the 5 algorithms across two cell types.

## 4 Conclusion

Altogether, this study sought to compare network inference methods and the impact of cyclic time as well as linear clock time on the accuracy of inferred networks. We found that timing-aware methods were essential to infer the gene regulatory networks associated with cell fate specification in the developing retina. The cyclic time of cell cycle had the most significant impact of the performance of network inference methods. New integrative metrics incorporating both cell cycle and linear time can account for both processes in regulatory network inference for developmental biology. Altogether, these results point to the need for extending regulatory network inference to cyclic processes and multi-scale temporal processes common to biological systems.

## Data and code availability

The mouse RPC data [1] can be accessed from https://www.ncbi.nlm.nih.gov/geo/query/acc.cgi?acc=GSE118614. The code that documented our findings is available at GitHub repo: https://github.com/aplazar1/WmB_2025_causal_GRN_inference.git.

## Acknowledgments

This material is based upon work supported by the National Science Foundation under Grant No. DMS-1929284 while the authors were in residence at the Institute for Computational and Experimental Research in Mathematics in Providence, RI, during the Women in Mathematical Computational Biology program held in January 2025. Additional funding was provided by R00NS122085, and Kavli Neuroscience Discovery Institute, and the Terkowitz Family Rising Professorship to G.L.S.-O. and NCI U24CA284156 to E.J.F. and G.L.S.-O.

